# Temporal Structure of Environmental Noise Controls the Localization and Tracking of Populations of Chemotactic Microorganisms

**DOI:** 10.64898/2026.05.07.723364

**Authors:** Gianna Arencibia, Martin Gutiérrez, Fivos Panetsos

## Abstract

Chemotactic microorganisms navigate fluctuating chemical landscapes by adjusting their motion in response to changes in locally sensed concentration. Consequently, population-level localization may depend not only on the magnitude of environmental fluctuations but also on their temporal structure. We investigate this dependence using a minimal agent-based model of noninteracting, Escherichia coli-inspired run-and-tumble swimmers in static and moving chemoattractant landscapes. We compare white, pink, Ornstein–Uhlenbeck, Brownian, and band-limited perturbations and examine how noise correlation time affects localization and tracking. Within the conditions studied, short-correlated fluctuations have relatively little effect, whereas slowly varying perturbations reduce retention, increase dispersion, and impair tracking. In the Ornstein–Uhlenbeck series, the strongest deterioration occurs when the noise correlation time approaches the characteristic run timescale. A low-frequency power fraction provides a useful empirical descriptor of these changes: localization data from different noise classes follow a similar trend when plotted against this quantity. Simulations with spatially localized advection further show that aggregate disruption depends on the placement of flow relative to the localization region. These findings link temporal noise structure and spatial transport to the robustness of population-level chemotactic organization in a minimal model.

## 1. Introduction

Motile microorganisms that sense and navigate chemical gradients—chemotaxis—provide a canonical example of active matter in which self-propelled agents process environmental information to generate directed motion^1^. In *Escherichia coli* and related species, chemotaxis relies on a finely tuned signaling cascade that couples chemoreceptor activity to the flagellar motors, allowing cells to bias run-and-tumble trajectories toward favorable conditions and away from harmful environments^2,3^. At the population level, this microscopic strategy underpins collective migration toward localized chemical cues and plays a central role in biofilm formation, host–pathogen interactions, and the spatial organization of microbial communities^4^.

Despite this detailed single-cell understanding, how microscopic sensing and motility rules translate into robust collective localization under realistic environmental conditions remains incompletely understood. In natural and physiological settings, chemotactic navigation rarely occurs in static or noise-free landscapes^5^. Instead, bacterial populations experience chemical signals that evolve in space and time, are continuously reshaped by diffusion, metabolism, and environmental flows, and exhibit fluctuations over a broad range of temporal scales^5,6^. These fluctuations introduce competing mechanisms chemotactic guidance, diffusive dispersion, and advective transport that jointly determine whether populations successfully localize, track moving targets, or fail to reach regions of interest^5-7^.

From a theoretical perspective, continuum models of Keller–Segel type and related stochastic frameworks have clarified how chemotactic populations aggregate, form waves, and approach the limits of sensing precision in the presence of molecular shot noise and receptor-level fluctuations^6-9^. However, most studies that incorporate noise treat it as temporally uncorrelated, effectively assuming white or rapidly decorrelating fluctuations in the chemical field or in intracellular signaling^8-10^. As a result, the role of temporal noise correlations in controlling population-level robustness has remained largely unexplored, even though many natural environments exhibit slowly varying or intermittently correlated chemical disturbances^5,6^.

Here we focus on this underexplored problem, namely on how temporal correlations in signal noise dictate chemotactic robustness. Specifically, we ask how the temporal structure of environmental fluctuations not just their amplitude controls the ability of chemotactic populations to localize and track targets. To address this question, we develop a minimal agent-based framework in which individually non-interacting run-and-tumble cells implement a temporal sensing strategy inspired by *E. coli* and respond to externally prescribed chemoattractant fields under confinement.^3^ Within this model, we systematically expose the population to static and moving signals perturbed by distinct classes of temporal noise, including white noise, pink noise, and correlated random motion, thereby spanning regimes from short-correlation, high-frequency fluctuations to slowly varying, long-correlation disturbances.

Our central message is that collective chemotactic targeting is a temporal filter: populations suppress high-frequency noise but collapse under slowly varying, correlated perturbations, defining intrinsic limits to programmable guidance. Using the existing simulations of white, pink, and correlated noise, we show that the temporal correlation structure of signal fluctuations, rather than noise amplitude alone, controls the transition from robust targeting to failure, both for static localization and for tracking of moving targets. We identify a critical correlation timescale comparable to the cellular response time beyond which chemotactic aggregates become unstable and tracking accuracy deteriorates sharply. This behavior connects conceptually to recent work on precision limits and information-theoretic constraints in single-cell chemotaxis, but, to our knowledge, no previous study has systematically mapped how different regimes of temporal noise (short-versus long-correlation) shape collective localization and target tracking at the population level^8-11^. Finally, we place these findings in the broader context of programmable microbial systems. In engineered or microfluidic environments, signal landscapes and flow conditions can be designed, yet they inevitably contain temporal variability^12^. Our results suggest that the temporal spectrum of these fluctuations is a key design parameter: by constraining low-frequency noise or by matching cellular response times to the characteristic fluctuation scales, one can enhance robustness of chemotactic targeting. Conversely, environments dominated by slowly varying, correlated perturbations may fundamentally limit the controllability of chemotactic collectives, even when average gradients are favorable. This perspective points toward strategies in which chemically programmable landscapes and controlled noise statistics are combined with physical stabilization mechanisms to harness chemotactic populations as adaptive, self-organizing agents in living materials, targeted delivery, and spatially structured biosystems^12,13^.

In summary, our work makes three main contributions to the physics of chemotactic active matter. First, we introduce temporal noise correlations specifically, noise correlation time and low-frequency spectral content as explicit control parameters for collective chemotactic robustness, going beyond prior studies that focused primarily on noise amplitude or assumed temporally uncorrelated fluctuations. Second, by systematically varying the correlation time of environmental perturbations, we identify a critical timescale, comparable to the mean run duration, that marks a sharp transition between robust collective localization and the breakdown of tracking and aggregation. Third, we establish a quantitative link between the low-frequency components of environmental noise spectra and the success or failure of chemotactic targeting, showing that slow fluctuations, rather than total variance, are the dominant predictors of collapse. Together, these advances reveal collective temporal filtering as a distinctive emergent property of chemotactic populations and provide a minimal quantitative framework for analyzing and designing robust chemotactic guidance in fluctuating environments.

## 2. Materials and Methods

### 2.1 Simulation Platform

All simulations were performed using GRO, an open-source stochastic agent-based framework for modeling spatially distributed microbial populations ^14^, and complemented by independent implementations in MATLAB (MathWorks, Natick, MA, USA). In both frameworks, individual bacteria are represented as autonomous agents that continuously sample the local chemoattractant concentration and update their motility based on internal decision rules. The simulation domain consists of a two-dimensional square region of size 600 × 600 μm, coupled to a diffusible chemoattractant field *S*(x, *t*)defined on a regular Cartesian lattice with spatial resolution Δ*x*.

The temporal evolution of the chemical field is governed by a reaction–diffusion equation of the form:

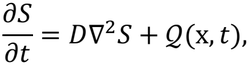

where *D*is the diffusion coefficient and *Q* (x, *t*)represents localized source terms corresponding to attractant emitters. The equation is integrated numerically using an explicit finite-difference scheme. Numerical stability was ensured by selecting Δ*t* and Δ*x* such that the Courant–Friedrichs–Lewy (CFL) condition for diffusion-dominated dynamics is satisfied. At each time step, agents sample the chemoattractant concentration at their instantaneous position by bilinear interpolation of the four nearest lattice nodes, since agent coordinates are not, in general, coincidence with the grid points on which the chemical field is defined.

Boundary conditions were adapted to the experimental configuration. In baseline scenarios, reflective (no-flux) boundary conditions were imposed on both agents and the chemical field. Under flow conditions, open boundary segments were introduced to model advective loss: agents crossing these boundaries were removed from the system, while the chemical field remained governed by no-flux conditions. The GRO framework was extended with custom modules implementing: (i) temporal chemotactic sensing, (ii) time-dependent signal sources, (iii) stochastic perturbations with controlled temporal correlations, and (iv) externally imposed flow fields (see Section X.X). Equivalent modules were implemented in MATLAB to reproduce the same dynamical rules and environmental conditions. Cross-validation between the two implementations confirmed that all key observables (localization efficiency, tracking error, and spatial dispersion) were consistent within statistical uncertainty.

Simulations were integrated with a time step of Δ*t* = 0.05 sand run for up to 10,000 steps (500 s). Numerical convergence was verified by halving the time step, resulting in changes below 2% in all reported observables. Simulation outputs were post-processed using MATLAB for trajectory analysis, statistical evaluation, and spectral decomposition of signal fluctuations. Each condition was repeated over n = 8 independent realizations with different random seeds to account for stochastic variability. Reported values correspond to ensemble averages, and variability is quantified using the standard error of the mean (SEM).

### 2.2 Agent-Based Chemotactic Model

Bacterial motility was modeled using an agent-based framework inspired by the canonical chemotactic behavior of *Escherichia coli*. Each cell is represented as a self-propelled particle in a two-dimensional domain, characterized by its position **r**_*i*_(*t*) = (*x*_*i*_, *y*_*i*_)and orientation *θ*_*i*_ (*t*). During run phases, cells move at constant speed *v*_0_along their instantaneous orientation. The propulsion speed was set to

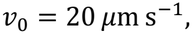

consistent with experimentally reported swimming speeds for *Escherichia coli* (typically 15–30 μm s^−1^), ensuring that the simulated dynamics operate within biologically realistic regimes. In discrete time, positions are updated as

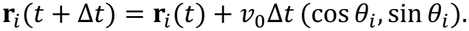

Motility follows a stochastic run-and-tumble dynamic in which cells alternate between: (i) runs, characterized by persistent motion, and (ii) tumbles, during which cells undergo rapid reorientation. Tumbles are modeled as instantaneous events that reset the orientation *θ*_*i*_ by drawing a new angle from a uniform distribution in [0,2*π*).

Transitions between run and tumble states are governed by a Poisson process with a time-dependent tumbling rate *λ*(*t*), modulated by the local chemoattractant signal. In the absence of chemotactic bias, the baseline tumbling rate is defined as

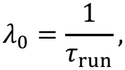

where the mean run duration was set to

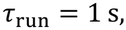

consistent with experimental measurements for *E. coli*. In discrete time, the probability of a tumble event within a time step Δ*t*is given by

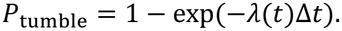

### Chemotactic modulation of tumbling rate

Chemotactic bias was implemented through a temporal sensing mechanism in which cells respond to changes in the chemoattractant concentration sampled along their trajectories. Let

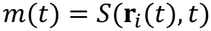

denote the local signal experienced by a cell. The chemotactic response is based on the temporal increment

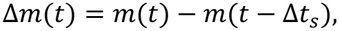

where Δ*t*_*s*_ is the sensing timescale. Importantly, the tumbling rate was modeled as a continuous function of the temporal signal increment, rather than a purely sign-based rule. Specifically, we used an exponential modulation:

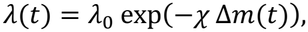

where *χ*is a sensitivity parameter controlling the strength of the chemotactic response. This formulation ensures that the tumbling rate responds not only to the sign but also to the magnitude of the sensed gradient. Positive temporal gradients (Δ*m* > 0) lead to an exponential suppression of tumbling, resulting in longer runs and stronger directional persistence. Conversely, negative gradients increase the tumbling rate, promoting reorientation.

### Rationale and scaling behavior

This continuous formulation resolves a key limitation of sign-based modulation rules, in which the tumbling rate depends only on the sign of Δ*m*and is therefore invariant to the amplitude of the chemoattractant field. In contrast, the exponential response introduces a scale-dependent chemotactic bias, such that stronger gradients produce proportionally stronger suppression of tumbling. This is essential to capture realistic behavior in which the aggregation dynamics depend on signal strength. From a biophysical perspective, this choice is consistent with experimental and theoretical descriptions of bacterial chemotaxis, where intracellular signaling pathways effectively map receptor occupancy changes into nonlinear modulation of motor switching rates. The exponential form provides a minimal phenomenological representation of this nonlinear transduction while preserving numerical stability and positivity of *λ*(*t*).

### Numerical constraints

To ensure physical consistency, the parameter *χ*was chosen such that: *λ*(*t*) > 0for all Δ*m*, variations in *λ*(*t*) remain within biologically plausible ranges, and the characteristic response remains comparable to the baseline tumbling rate *λ*_0_. The stochasticity of the run-and-tumble process introduces intrinsic variability in cell trajectories, capturing the combined effects of intracellular signaling noise, flagellar motor fluctuations, and environmental heterogeneity. No explicit rotational diffusion term is included; instead, directional randomness arises solely from tumble events and probabilistic switching dynamics. Importantly, no direct inter-agent interactions (e.g., steric exclusion, alignment, or hydrodynamic coupling) are incorporated. As a result, all emergent collective behaviors arise exclusively from the coupling between individual chemotactic responses and the spatiotemporal structure of the external chemoattractant field. This minimal formulation enables isolation of the role of signal topology, temporal variability, and external perturbations in shaping population-level organization.

### 2.3 Signal Modeling and Chemotactic Capture Region

The chemical environment was modeled as a spatially and temporally varying chemoattractant field *S*(**x**, *t*), which defines the external cues guiding bacterial motion. Rather than restricting the analysis to a single signal configuration, we systematically varied the spatiotemporal properties of *S* to investigate how different classes of environmental stimuli shape collective chemotactic behavior.

The chemoattractant field was constructed as the superposition of localized sources:

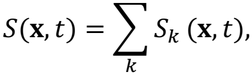

where each source *S*_*k*_ generates a spatial gradient centered at position **x**_*k*_ (*t*). Unless otherwise specified, individual sources were modeled as smooth, radially decaying profiles (e.g., Gaussian-like fields), ensuring well-defined gradients over a characteristic spatial scale comparable to the aggregation region.

To disentangle the effects of spatial structure, temporal dynamics, and environmental variability, we considered three progressively complex signal regimes: static fields, dynamic fields, and stochastically perturbed fields.

To formalize the spatial domain within which chemotactic guidance remains effective, we define the chemotactic capture region Ω_*c*_ as the set of positions where chemotactic drift can overcome stochastic reorientation and external perturbations:

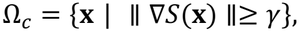

where ∥ ∇*S*(**x**) ∥ is the local magnitude of the chemoattractant gradient and *γ* is an effective threshold that represents the minimal gradient required to produce a statistically significant bias in run duration relative to stochastic tumbling. This threshold implicitly depends on the intrinsic properties of the agents, including the mean run duration *τ*_run_, the chemotactic sensitivity parameters *α, β*, and the noise level in the sensed signal. Within Ω_*c*_, chemotactic drift dominates and cells can maintain directed motion toward the signal source. Outside this region, stochastic reorientation and external perturbations dominate, leading to effectively diffusive or advectively driven motion^15-17^.

### Interpretation and coupling to flow and noise

The capture region provides a unifying framework to interpret the effects of temporal noise and advective transport: temporal noise, when fluctuations occur on timescales *τ*_0_ ≫ *τ*_run_, the effective position of the signal drifts, causing cells to exit Ω_*c*_and lose localization, and advective transport, when cells are displaced outside Ω_*c*_, chemotactic gradients are no longer sufficient to restore directed motion, leading to aggregate destabilization. Thus, the stability of chemotactic aggregates is governed by whether environmental perturbations—temporal or spatial—displace agents beyond the capture region^17-19^.

### Characteristic length scale of the capture region

For analytical interpretation, the chemotactic capture region can be associated with a characteristic spatial scale *R*_*c*_, defined as the distance from the signal center at which the gradient magnitude falls below the threshold *γ*:

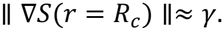

For radially symmetric signal profiles with characteristic amplitude *S*_0_, this condition yields an estimate of the capture radius:

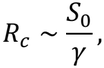

up to a geometry-dependent prefactor that depends on the specific functional form of the signal field. This scaling reflects the balance between signal strength and the minimal gradient required to bias chemotactic motion. Higher signal amplitudes *S*_0_ extend the spatial region over which gradients remain detectable, increasing *R*_*c*_, whereas larger thresholds *γ*—associated with weaker sensitivity or stronger stochasticity—reduce the effective capture region^16,17^.

### 2.4 Signal Fields

#### Static Signal Fields

In the baseline configuration, signal sources were fixed in space, 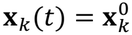, generating time-invariant concentration landscapes. Multiple sources were introduced to create overlapping gradients and spatially structured attraction basins. Under these conditions, the chemoattractant field defines a static landscape in which bacterial dynamics are governed solely by individual chemotactic responses. The characteristic gradient magnitude scales as ∇*S* ∼ *S*/*σ*_*s*_, setting the strength of directional bias experienced by the agents. This regime provides a reference scenario for quantifying aggregation efficiency, spatial precision, and steady-state population structure in the absence of temporal variability. The static case serves two main purposes: (i) to identify configurations that maximize or hinder localization, and (ii) to define benchmark “best-case” and “worst-case” conditions used in subsequent perturbation experiments^15,16^.

#### Dynamic Signal Fields

To model non-stationary environments, signal sources were allowed to evolve in time, such that **x**_*k*_ = **x**_*k*_ (*t*). This generates moving concentration landscapes that require continuous adaptation of bacterial trajectories. Source trajectories were prescribed and parameterized by their velocity *v*_*f*_ and spatial pattern (e.g., sinusoidal or translational motion), enabling controlled exploration of different dynamical regimes. In this setting, the chemotactic task shifts from static localization to dynamic tracking. The interplay between signal velocity and cellular response time defines a natural limit to tracking performance. When the characteristic displacement over one run length *v*_*f*_ *τ*_run_becomes comparable to or larger than the gradient length scale *σ*_*s*_, cells are unable to reliably infer the direction of motion. This results in increased tracking errors, delayed response, and partial fragmentation at the population level^17-19^.

#### Stochastic Signal Perturbations

To incorporate environmental variability, stochastic fluctuations were introduced into the signal field. Perturbations were applied either to the source position **x**_*k*_(*t*)or to the source amplitude *A*_*k*_ (*t*), generating time-dependent variability in the chemoattractant landscape. Three classes of noise were considered, each representing distinct physical and biological sources of variability with well-defined temporal structure: White noise (uncorrelated, high-frequency fluctuations), Pink noise (1/f) (long-range temporally correlated fluctuations), and Correlated stochastic motion (continuous random trajectories of signal sources). These noise models are not arbitrary abstractions, but correspond to different classes of environmental variability encountered by chemotactic microorganisms.

White noise represents fast, uncorrelated fluctuations arising from molecular-scale processes such as stochastic ligand binding and unbinding at chemoreceptors, rapid diffusion-driven concentration fluctuations, and local turbulence at microscales. These fluctuations occur on timescales much shorter than the bacterial response time and are therefore expected to be partially averaged out by temporal sensing mechanisms^20,23^.

In contrast, pink noise (1/f) captures scale-invariant, long-memory fluctuations commonly observed in biological and environmental systems. Such noise emerges from processes with broad temporal spectra, including metabolic activity fluctuations, intermittent secretion of signaling molecules, and heterogeneous diffusion in complex media such as biofilms or porous environments. These fluctuations introduce persistent low-frequency components that evolve on timescales comparable to or longer than chemotactic adaptation, making them particularly relevant for testing the limits of temporal sensing^19,20^.

Finally, correlated stochastic motion models smooth, continuous displacement of signal sources, representing situations in which chemoattractant fields are advected or reshaped by underlying physical processes. Examples include nutrient plumes transported by fluid flow, moving signaling fronts in microbial communities, or dynamically evolving chemical hotspots in tissues. Unlike purely stochastic intensity fluctuations, this regime introduces structured, directional variability in the signal landscape, directly challenging the ability of cells to infer gradient direction over time^19,21^.

By combining these three noise classes, we span a broad spectrum of temporal variability, from fast, memoryless perturbations to slow, correlated environmental changes and structured signal motion. This allows us to isolate how the temporal statistics of fluctuations—rather than their amplitude alone— control the robustness of chemotactic targeting.

#### Stochastic processes based on exponentially correlated noise

To systematically control temporal correlations, we further introduced a continuous family of stochastic processes based on exponentially correlated noise. Specifically, fluctuations were generated using an Ornstein–Uhlenbeck process:

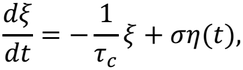

where *τ*_0_is the correlation time, *σ* sets the noise amplitude, and *η*(*t*)is Gaussian white noise. In this formulation, the limit *τ*_*c*_ → 0recovers uncorrelated noise, while increasing *τ*_*c*_ produces progressively slower and more persistent fluctuations. By sweeping *τ*_0_across a range spanning values much smaller and much larger than the characteristic run duration *τ*_run_, we constructed a continuum of noise regimes that interpolate between rapidly fluctuating and slowly varying environments. The Ornstein–Uhlenbeck process was employed as a minimal and physically grounded model of temporally correlated fluctuations. Unlike white noise, which assumes instantaneous decorrelation, the Ornstein–Uhlenbeck process introduces a finite correlation timescale *τ*_*c*_, enabling controlled interpolation between rapidly fluctuating and slowly varying environments.

This choice is motivated by the fact that many biological and physical processes governing chemoattractant dynamics—such as diffusion with degradation, metabolic production, and transport in heterogeneous media—naturally generate fluctuations with finite temporal memory and approximately exponential autocorrelation structure. In addition, intracellular chemotactic signaling networks exhibit characteristic response and adaptation timescales, implying that cells effectively filter environmental signals over finite temporal windows. As a result, the relative magnitude of *τ*_*c*_ with respect to the mean run duration *τ*_run_ is expected to play a central role in determining chemotactic performance.

By systematically varying *τ*_0_, the Ornstein–Uhlenbeck framework enables a direct exploration of how temporal correlations in environmental signals interact with the intrinsic response timescale of chemotactic cells, providing a mechanistic basis for identifying robustness limits and failure regimes in collective targeting. Importantly, stochastic perturbations were applied selectively to representative signal configurations identified in the static and dynamic regimes. This targeted approach allows direct comparison of robustness across noise types while avoiding redundant sampling of parameter space^20-23^.

#### Signal Fields as Control Parameters for Emergent Behavior

Across all regimes, the chemoattractant field *S*(**x**, *t*)acts as the primary external control parameter governing system behavior. Its key characteristics—gradient magnitude, spatial extent *σ*_*s*_, temporal dynamics *v*_*f*_, and noise correlation time *τ*_*c*_—define a reduced but expressive parameter space controlling chemotactic performance. Within this framework, population-level organization emerges from the interplay between local temporal sensing and global signal structure. In particular, collective behavior is determined by whether environmental variations occur within or outside the effective sensing bandwidth of the agents. This formulation enables a systematic investigation of how programmable chemical environments—characterized by their spatial topology, temporal variability, and noise spectra—control the formation, stability, and reconfiguration of collective behaviors in active biological systems^24-27^.

### 2.5 Flow Field Modeling and Coupling to Chemotactic Dynamics

To investigate the impact of spatially heterogeneous transport on chemotactic organization, externally imposed flow fields were introduced as localized advective perturbations acting on bacterial trajectories. These perturbations were applied after the system reached a quasi-stationary aggregated state under static signal conditions, enabling isolation of the effect of flow on aggregate stability rather than formation dynamics. The motion of each agent results from the superposition of intrinsic self-propulsion and externally imposed advection:

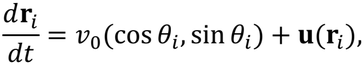

where *v*_0_is the swimming speed and **u**(**r**)is the local flow velocity^25,28^.

#### Spatial structure and geometry of the flow field

The flow field was defined as one or two spatially confined vertical strips located at the lateral boundaries of the domain. Within these regions, flow was oriented along the vertical direction (bottom-to-top), mimicking localized transport streams such as those encountered in microfluidic channels or porous media. Formally, the velocity field was defined as:

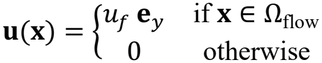

where *u*_*f*_ is the flow intensity and Ω_flow_ denotes the spatial region occupied by the flow strips. Each strip is characterized by three geometric parameters: Inner-edge position *d*, the distance between the inner boundary of the strip and the signal center; Width *W*, the spatial extent of the strip along the transverse direction; and Topology, either unilateral (left or right) or bilateral (both sides). Importantly, the strip was always defined such that its inner boundary is anchored to a given iso-concentration contour of the signal field, and the width extends outward from this boundary. This ensures a controlled and physically meaningful parametrization of the flow–signal interaction region^29-31^.

#### Coupling between advection and chemotactic sensing

The flow field does not directly modify the chemotactic decision rules, but alters the trajectories along which cells sample the chemoattractant field:

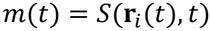

Therefore, advection perturbs the temporal signal increments Δ*m*(*t*), effectively biasing or disrupting the inferred gradient direction. This induces competition between chemotactic drift, arising from biased run-and-tumble dynamics, and advective transport, which displaces cells independently of their orientation. The balance between these mechanisms determines whether cells remain within the region where gradients can guide reorientation^24,26,28^.

#### Effective capture region and spatial overlap

A central concept emerging from this formulation is the existence of an effective chemotactic capture region, defined as the spatial domain in which signal gradients are sufficiently strong to retain cells against stochastic reorientation and external perturbations. Flow-induced destabilization occurs when the advective field overlaps with this region. When the flow is positioned outside this domain, chemotactic drift dominates and aggregation remains stable. Conversely, when the flow intersects the capture region, advection removes cells faster than they can reorient and return, leading to aggregate depletion. Consistent with the simulation results, this defines a finite spatial range of influence of the flow, on the order of the characteristic signal length scale (≈80–140 μm), beyond which the perturbation becomes negligible^28,32^.

#### Topological effects and asymmetry

The spatial topology of the flow field introduces qualitatively distinct regimes of aggregate response. Unilateral flow induces partial depletion while preserving a residual aggregate, as cells can re-enter the signal region from the unperturbed side. In contrast, bilateral flow generates a divergent advective field that removes cells from both flanks, suppressing recovery pathways and leading to aggregate collapse. Asymmetric configurations produce intermediate behavior, creating protected regions with reduced transport that enable partial recovery and enhanced retention. These results demonstrate that aggregate stability depends not only on flow intensity, but critically on the availability of spatial pathways that allow reinjection of cells into the signal region^29,31,33^.

#### Dominant geometric control parameters

This formulation allows disentangling the relative contributions of flow geometry to aggregate stability. The position of the inner edge, *d*, emerges as the primary control parameter, as it determines whether the flow overlaps with the high-gradient core of the signal. Once this region is intersected, destabilization is triggered. In contrast, the strip width, *W*, plays a secondary role, modulating the extent of perturbation without significantly altering the onset of instability. In asymmetric configurations, the total advective input dominates over its spatial partitioning, except in cases where asymmetry generates protected regions that sustain local retention and enable recovery. This hierarchy of control parameters is reflected in the nonlinear system response, including sharp destabilization thresholds as the flow approaches the signal core^27,33^.

#### Biophysical relevance

The imposed flow can be interpreted as a generic external perturbation acting on the chemotactic signal landscape, effectively filtering and reshaping the collective behavior of the population. Such perturbations are common in realistic environments, including natural fluid systems and engineered settings. The adopted representation captures key features of transport in physically relevant scenarios, such as microfluidic devices with lateral injection streams and transport through heterogeneous porous media. In these systems, flow is typically spatially localized and anisotropic rather than uniform, making strip-based perturbations a minimal yet physically grounded framework to study flow– chemotaxis coupling^29,31^.

### 2.6 Experimental Design

The simulation study was structured as a sequence of progressively complex experimental scenarios designed to isolate and quantify the impact of key environmental factors—namely spatial signal structure, temporal dynamics, stochastic variability, and external transport—on collective chemotactic behavior. This hierarchical design enables a controlled transition from idealized conditions to increasingly realistic environments while preserving mechanistic interpretability. In all experiments, bacterial populations were initialized with uniformly random positions and orientations. Each condition was evaluated across multiple independent realizations to ensure statistical robustness and reproducibility of observed behaviors.

#### Static Signal Localization

In the baseline scenario, agents were exposed to time-invariant chemoattractant fields defined by fixed spatial sources. Multiple signal configurations and intensity profiles were explored to characterize the ability of the population to aggregate and maintain stable localization. The primary observable in this regime was the localization efficiency, defined as the fraction of agents within a target region centered at the signal source. This scenario establishes reference conditions for optimal and suboptimal aggregation, which are subsequently used as benchmarks for evaluating robustness under dynamic and stochastic perturbations.

#### Dynamic Signal Tracking

To probe adaptive behavior in non-stationary environments, signal sources were allowed to evolve over time following prescribed trajectories **x**_*f*_(*t*), parameterized by their velocity *v*_*f*_ and spatial pattern. In this regime, the task shifts from static localization to continuous tracking of a moving target. Performance was quantified using two primary metrics: the mean tracking error, defined as the average distance between agents and the instantaneous signal position, and the response delay, capturing the temporal lag between signal displacement and population adaptation. By systematically varying *v*_*f*_, this phase characterizes the dynamic limits of collective chemotactic tracking and identifies regimes of coherent following, partial fragmentation, and tracking failure.

#### Signal Uncertainty and Noise

To assess robustness under environmental variability, stochastic perturbations were introduced into the signal field. Rather than exhaustingly sampling the full parameter space, noise was selectively applied to representative configurations identified in previous phases, specifically those exhibiting high and low localization performance. Different noise regimes—white, pink, and temporally correlated stochastic motion—were imposed on either the position or intensity of signal sources. This design enables direct comparison of how distinct temporal structures of environmental fluctuations affect aggregation stability, spatial dispersion, and tracking performance. By focusing on extreme reference cases, this approach maximizes interpretability while minimizing redundancy, allowing clear identification of conditions under which collective behavior remains robust or breaks down.

#### Correlation-Time Sweep and Spectral Analysis

To systematically quantify the role of temporal correlations, we performed an additional set of simulations in which the noise correlation time *τ*_*c*_ was varied over several orders of magnitude. Starting from representative static and dynamic configurations, exponentially correlated perturbations were imposed with *τ*_0_spanning regimes from *τ*_*c*_ ≪ *τ*_run_ to *τ*_*c*_ ≫ *τ*_run_. For each condition, we computed localization efficiency, spatial dispersion, persistence time of aggregated states, and tracking error. This design enables mapping of transitions between robust and failed chemotactic targeting as a function of *τ*_*c*_, while minimizing redundant sampling of parameter space. To relate environmental fluctuations to the intrinsic sensing capabilities of chemotactic agents, we quantified the low-frequency content of the signal using the power spectral density *S*(*f*). The low-frequency power,

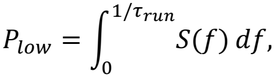

was defined as the integrated spectral energy over frequencies lower than the inverse of the mean run duration *τ*_*run*_. This threshold reflects the intrinsic temporal resolution of chemotactic sensing, as fluctuations occurring on timescales shorter than *τ*_*run*_ are effectively averaged out, whereas slower fluctuations directly influence directional decisions. Therefore, *P*_*low*_ provides a physically grounded measure of the fraction of environmental variability that is dynamically relevant for chemotactic navigation.

#### Flow-Induced Perturbations

Finally, the stability of emergent population states was evaluated under externally imposed flow conditions. To isolate the effect of transport on already formed structures, flow fields were introduced after the system reached a quasi-stationary aggregated configuration, thereby probing the resilience of established aggregates rather than their formation dynamics. The flow was implemented as a spatially structured velocity field **u**(**x**), superimposed on the intrinsic self-propelled motion of individual agents. Consistent with the flow model described in Section 2.3.5, advection was applied through localized vertical strips positioned along the left and/or right boundaries of the domain, with flow directed along the *y*-axis (bottom-to-top). This configuration enables independent control of flow topology, spatial extent, and geometric overlap with the chemoattractant field.

The following parameters were systematically varied: Flow intensity *u*_*f*_, magnitude of the advective velocity, Topology: unilateral (single-sided) versus bilateral (two-sided) flow; Geometric configuration, strip width *W*and distance *d*between the inner edge of the flow region and the signal center; and Asymmetry, independent variation of velocity and width on opposing sides (*u*_*L*_ ≠ *u*_*R*_, *W*_*L*_≠ *W*_*R*_).

Within each flow region, the velocity field was spatially uniform and time-independent, generating a controlled vertical advective flux. This design enables systematic exploration of the competition between: chemotactic drift, arising from biased run-and-tumble dynamics, and advective transport, which displaces agents independently of local sensing

### Central hypothesis tested in this phase

The central hypothesis tested in this phase is that aggregate stability is governed not only by the magnitude of advective forcing, but by its spatial overlap with the effective chemotactic capture region. By varying the position and extent of flow relative to the signal landscape, we quantify the conditions under which chemotactic attraction can compensate for external transport, or conversely, is overcome by it. System response was quantified in terms of: structural persistence, via sustained localization efficiency; aggregate depletion, measured as the loss of agents within the focal region; and recovery capacity, defined as the ability of displaced agents to re-enter the signal region.

This framework defines a controlled and physically interpretable parameter space for investigating how spatially heterogeneous flow fields modulate the stability, resilience, and breakdown of chemotactic aggregates.

### 2.7 Quantitative Metrics

Simulation outputs were analyzed using custom MATLAB scripts. To quantitatively characterize the collective behavior of chemotactic populations across different signal regimes, a set of metrics capturing localization, tracking performance, spatial organization, and robustness was defined. All quantities were computed from agent positions **r**_*i*_(*t*)and, when applicable, the instantaneous position of the signal source **x**_*f*_(*t*). Reported values correspond to mean ± standard deviation across independent simulation replicates.

#### Localization Efficiency

Localization efficiency was defined as the fraction of agents contained within a circular target region of radius *R*centered at the signal source:

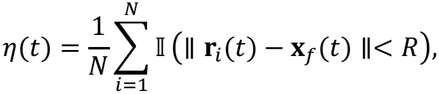

where *N*is the total number of agents and I(⋅) is the indicator function. The radius *R* was defined in relation to the chemotactic capture region introduced in Section 2.3. Specifically, *R* was chosen to be proportional to the characteristic capture radius *R*_*c*_, such that:

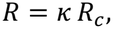

with *κ* ∼ *O*(1) a dimensionless factor ensuring that the target region encompasses the spatial domain over which chemotactic drift effectively counterbalances stochastic reorientation and external perturbations. This choice ensures that *η*(*t*)provides a physically grounded measure of aggregation, quantifying the fraction of agents that remain within the dynamically relevant sensing region of the signal. As a result, variations in *η*(*t*)directly reflect changes in the ability of the population to remain confined within the effective chemotactic capture region under different environmental conditions.

#### Tracking Error

For dynamic signal conditions, tracking performance was quantified through the mean tracking error:

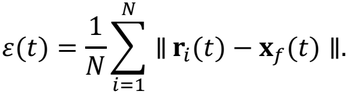

This metric measures the average spatial deviation between agents and the moving signal source, providing a direct estimate of tracking accuracy at the population level. To quantify variability across independent realizations and enable statistical comparison between conditions, we computed the standard error of the mean (SEM) of the tracking error:

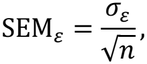

where *σ*_*ε*_ is the standard deviation of the tracking error across *n* independent simulation replicates. The SEM provides an estimate of the uncertainty associated with the sample means and was used to construct error bars and perform statistical comparisons in the Results section.

#### Spatial Dispersion

Population spread was quantified using the second central moment of the agent distribution:

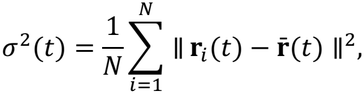

where

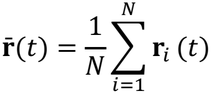

is the population centroid. This metric captures the spatial extent of the population and distinguishes between compact aggregates and dispersed configurations, being particularly sensitive to noise-induced fragmentation and loss of coherence. To quantify variability across independent realizations, we computed the standard error of the mean (SEM) of the spatial dispersion:

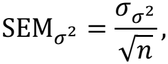

where 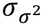 is the standard deviation of *σ*^2^ across *n*independent simulation replicates. This quantity provides an estimate of the uncertainty in the ensemble-averaged dispersion and was used for statistical comparison between experimental conditions.

#### Persistence Time

The temporal stability of collective organization was characterized by the persistence time *τ*_*p*_, defined as the longest continuous time interval during which the localization efficiency remains above a predefined threshold:

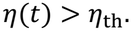

The threshold *η*_th_ was selected based on baseline aggregation levels in static conditions, ensuring that *τ*_*p*_ reflects meaningful retention of collective structure rather than transient fluctuations.

#### Success Probability

To quantify robustness across stochastic realizations, we defined the success probability:

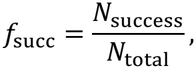

where *N*_success_ denotes the number of simulations achieving sustained localization (i.e., *η*(*t*) > *η*_th_ for a minimum persistence time), and *N*_total_ is the total number of realizations. This metric captures the reliability of chemotactic targeting under variability.

#### Temporal correlation and spectral metrics

To characterize the temporal structure of environmental fluctuations, we computed both time-domain and frequency-domain descriptors of the chemoattractant signal. The effective noise correlation time was defined as:

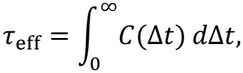

where *C*(Δ*t*)is the normalized temporal autocorrelation function of the signal sampled at fixed probe locations or at the population centroid, with *C*(0) = 1. This quantity provides a measure of the persistence of environmental fluctuations. In addition, the power spectral density *S*(*f*)of the signal was estimated using Welch’s method. To isolate fluctuations relevant to chemotactic sensing, we computed the low-frequency power:

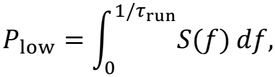

where *τ*_run_ is the mean run duration. This frequency threshold reflects the intrinsic temporal resolution of the chemotactic mechanism: fluctuations occurring on timescales shorter than *τ*_run_ are effectively averaged out by temporal sensing, whereas slower fluctuations directly influence directional decisions. By correlating *τ*_-eff_ nd *P*_low_ with localization efficiency, persistence time, and success probability across conditions, we assessed the extent to which temporal correlations and low-frequency spectral content predict the robustness or breakdown of collective chemotactic targeting. ^3^–^5^

#### Statistical Analysis

Statistical comparisons between experimental conditions were performed using one-way analysis of variance (ANOVA), which tests differences in the mean values of a given metric across multiple groups while controlling the overall type I error rate. The use of ANOVA is justified by the need to compare multiple environmental regimes (e.g., noise types, flow configurations, and signal velocities) and to identify global transitions between distinct behavioral states. When a significant effect was detected, pairwise comparisons were conducted using Tukey’s honestly significant difference (HSD) post-hoc test, which corrects for multiple comparisons and identifies statistically significant differences between specific conditions. Statistical significance was assessed at a threshold of *p* < 0.05. This analysis framework is appropriate for comparing ensemble-averaged quantities obtained from independent simulation replicates, where variability arises from stochastic dynamics and initial condition sampling.

## 3. Results

### 3.1 Static Signal Localization

We first establish a quantitative baseline by analyzing the collective behavior of chemotactic populations in static Gaussian chemoattractant fields of varying amplitude *S*_0_, with all other parameters held constant. This configuration isolates the intrinsic capacity of temporally sensing agents to generate spatial organization in the absence of environmental variability or external transport. As shown in Figure 1A, populations initialized from spatially homogeneous conditions spontaneously evolve toward localized aggregates centered at the signal maximum. The spatiotemporal snapshots (t = 0, 50, 150, 500 s) reveal a progressive collapse from a diffuse initial distribution to a compact, source-centered cluster. This transition occurs despite the absence of direct inter-agent interactions, confirming that aggregation arises solely from the coupling between individual temporal sensing and the spatial structure of the chemoattractant field. The dynamics exhibit a transition from an initially diffusive regime to a drift-dominated regime, in which biased run-and-tumble trajectories drive systematic convergence toward the maximum signal.

**Figure 1.**
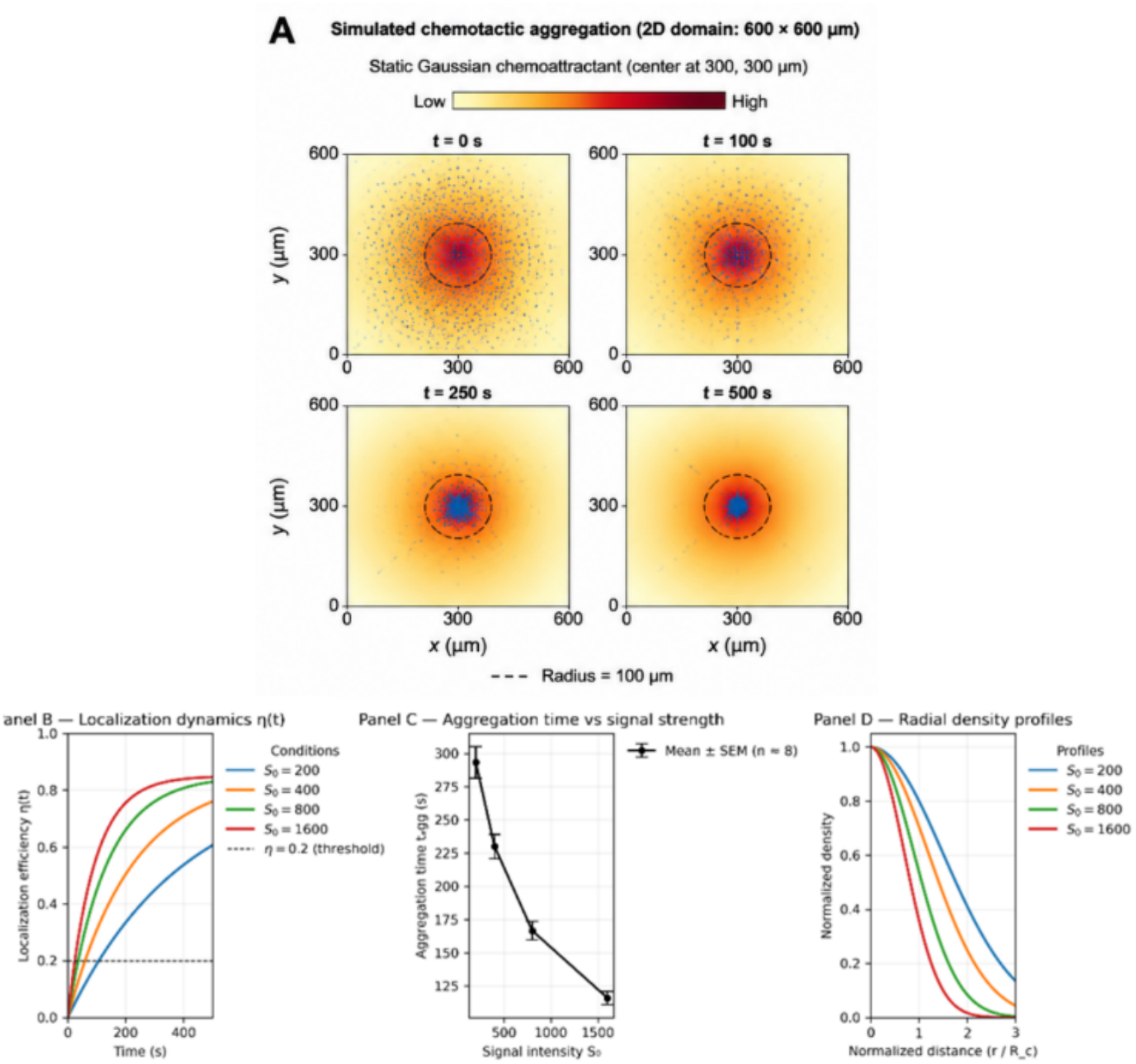
Chemotactic aggregation in static signal fields. (A) Spatiotemporal evolution of a bacterial population in a static Gaussian chemoattractant field, showing the transition from a homogeneous distribution to a localized aggregate within the capture region (dashed circle). (B) Localization efficiency η(t) for increasing signal amplitudes S_0_ illustrating faster convergence and higher steady-state localization at larger S_0_ (C) Aggregation time t_agg_ as a function of S_0_ (mean ± SEM), revealing a rapid decrease followed by saturation at high amplitudes. (D) Radial density profiles normalized by the capture radius R_c_ showing increasing spatial confinement of the population with signal strength.

The temporal evolution of localization efficiency, η(t), exhibits a strong dependence on signal amplitude *S*_0_(Figure 1B). At low signal intensities, η(t) remains close to baseline values, indicating that stochastic reorientation dominates over chemotactic bias. Quantitatively, for *S*_0_ = 200, the steady-state localization remains low (*η*_*ss*_ = 0.099), reflecting ineffective gradient sensing. Increasing *S*_0_ produces both faster convergence and higher steady-state localization, with *η*_.._rising monotonically to 0.949 at *S*_0_ = 5000(Table 1). Notably, signal amplitudes *S*_0_ ≤ 1000yield *η*_*ss*_< 0.25, defining an effective detection threshold below which chemotactic bias is insufficient to overcome diffusive dispersal. This threshold behavior is consistent with the gradual amplification of temporal signal increments governing the modulation of tumbling dynamics.

This dependence is further quantified by the aggregation time *t*_agg_, shown in Figure 1C. Above the detection threshold, *t*_agg_ decreases sharply with increasing *S*_0_, from 104.3 ± 11.4s at *S*_0_ = 1000to 8.5 ± 0.6s at *S*_0_ = 5000, before reaching a saturation regime. This behavior arises from the nonlinear modulation of the tumbling rate,

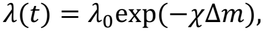

which amplifies differences in temporal signal increments. At low *S*_0_, small values of Δ*m*produce weak modulation of *λ*(*t*), resulting in dynamics close to unbiased diffusion. As *S*_0_ increases, larger temporal gradients strongly suppress tumbling, extending run durations and enhancing directional persistence. The saturation of *t*_agg_ at high *S*_0_, visible in Figure 1C, indicates that aggregation speed becomes limited by intrinsic dynamical constraints—such as finite swimming velocity and geometric confinement— rather than by sensing strength.

The spatial organization of the aggregated state is characterized by the radial density profiles shown in Figure 1D. For sufficiently large *S*_0_, the population becomes tightly confined within the chemotactic capture region. This is quantitatively reflected in the reduction of the aggregate half-width at half-maximum (HWHM), which decreases from 243.5 µm at low signal amplitude (*S*_0_= 100) to 33.0 µm at *S*_0_ = 5000, well within the capture radius *R* = *σ*_*s*_ = 100µm (Table 1). In contrast, weak signals produce broad spatial distributions extending far beyond *R*, indicating incomplete confinement. These results demonstrate that the capture region defines the effective spatial domain within which chemotactic drift overcomes stochastic reorientation and maintains localization.

Taken together, the four panels of Figure 1 provide a consistent multiscale picture of chemotactic aggregation in static environments. Panel A establishes the emergence of spatial organization, Panel B quantifies its temporal dynamics, Panel C identifies the scaling of the aggregation timescale, and Panel D reveals the spatial confinement mechanism. Crucially, these results establish the existence of a signal-dependent aggregation threshold and identify two emergent scales governing the process: a convergence timescale *t*_agg_(*S*_0_)controlled by tumbling-rate modulation, and a spatial confinement scale set by the capture radius *R*. These quantities define a well-characterized baseline regime for chemotactic organization and provide a reference framework for analyzing the effects of temporal variability, stochastic perturbations, and external transport in subsequent sections.

### 3.2 Dynamic Signal Tracking

Having established the static baseline, we next examined the ability of chemotactic populations to track a time-dependent signal, focusing on how tracking fidelity depends on the velocity of a moving chemoattractant source *v*_*f*_. In contrast to the static case, this configuration introduces a competition between the intrinsic response timescale of the population and the rate at which the environment changes. Representative spatial snapshots at equivalent phases of the source trajectory reveal distinct dynamical regimes as a function of *v*_*f*_ (Figure 2A).

**Figure 2.**
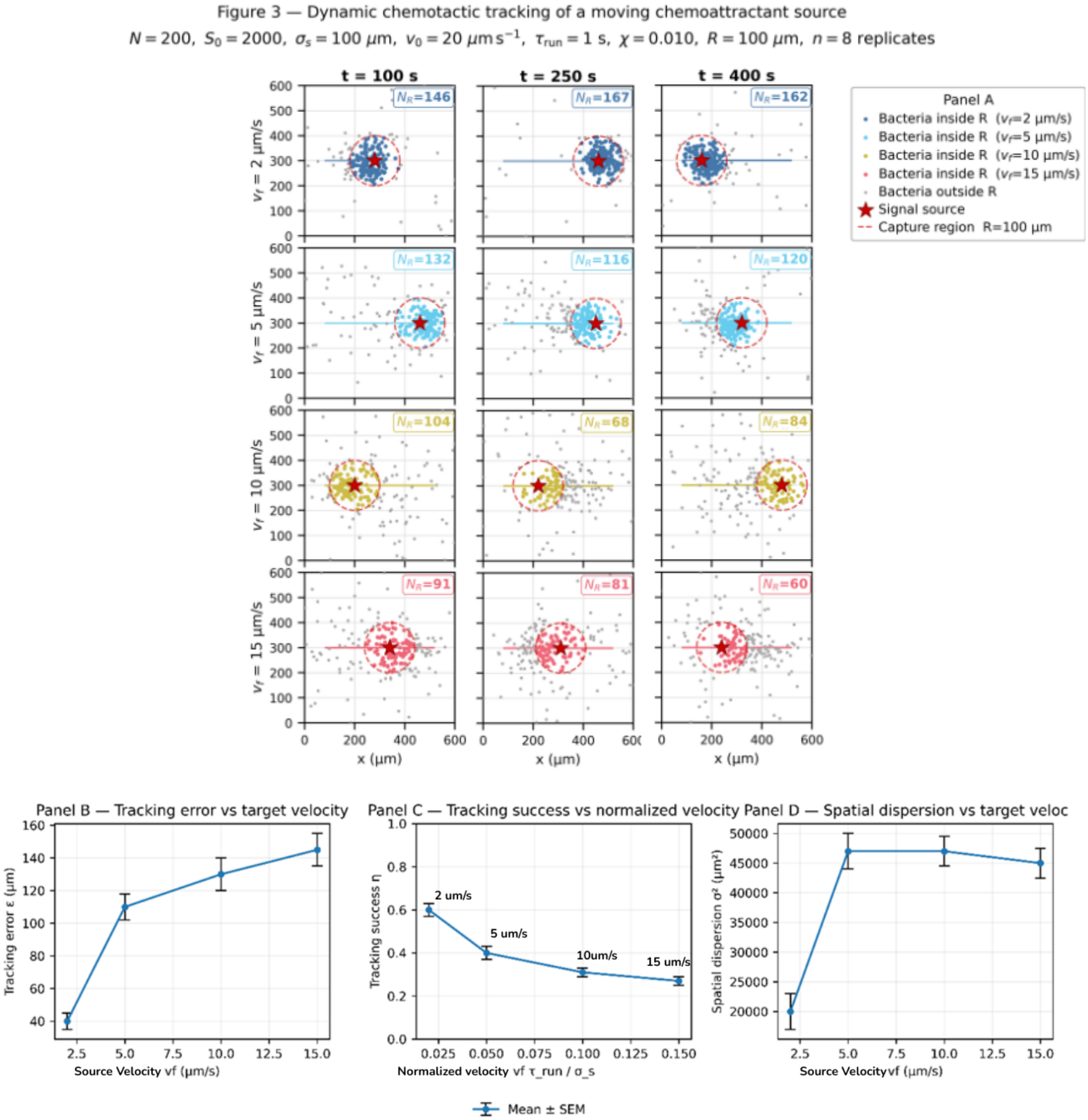
Dynamic chemotactic tracking of a moving chemoattractant source. (A) Spatial snapshots at different times for increasing source velocities v_f_ showing the transition from coherent tracking (low v_f_) to lag, fragmentation, and loss of co-localization at high v_f_. The dashed circle indicates the capture region. (B) Mean tracking error εincreases with v_f_ reflecting growing lag between the population centroid and the source. (C) Tracking success ηdecreases as a function of normalized velocity v_f_ τ_run_/σ_s_, revealing a transition to a degraded tracking regime. (D) Spatial dispersion σ^2^increases with v_f_, indicating progressive loss of aggregate coherence (mean ± SEM).

At low velocity (*v*_*f*_ = 2 *μ*m/s), the population remains coherently localized around the moving source across all time points, with the centroid maintaining a displacement comparable to or smaller than the capture radius *R*. This regime is characterized by effective dynamic tracking, in which the aggregate follows the source trajectory with a modest lag while preserving spatial coherence. At intermediate velocities (*v*_*f*_ = 5 *μ*m/s), the population begins to lag systematically behind the source. The centroid– source separation becomes comparable to the capture radius, and the aggregate exhibits increased spatial dispersion, indicating partial loss of confinement. Although tracking is still maintained, it occurs with reduced efficiency and increased elongation along the direction of motion. At higher velocities (*v*_*f*_ = 10 *μ*m/s), the system approaches a transition regime. The population centroid remains near or beyond the boundary of the capture region, and the aggregate becomes visibly fragmented. Subpopulations fail to remain localized near the moving maximum, indicating that the chemotactic response is no longer sufficient to reorganize the population on the timescale imposed by the source motion. Finally, at the highest velocity (*v*_*f*_ = 15 *μ*m/s), tracking breaks down completely. The centroid remains consistently displaced beyond the capture radius, and the population exhibits a diffuse and elongated structure with no stable co-localization with the source. This regime corresponds to a loss of dynamic tracking, where environmental changes outpace the intrinsic response of the system.

These qualitative observations are quantitatively captured by the dependence of tracking error *ε*, tracking success *η*, and spatial dispersion *σ*^2^on source velocity (Figure 2B–D). The mean steady-state tracking error increases monotonically with *v*_*f*_, from 73.0 ± 1.3 *μ*mat *v*_*f*_ = 2 *μ*m/sto 173.3 ± 1.1 *μ*mat *v*_*f*_ = 15 *μ*m/s, reflecting the growing lag between the population centroid and the moving source (Figure 2B). In parallel, tracking success decreases from *η* = 0.829 ± 0.005to 0.281 ± 0.002, indicating that an increasingly smaller fraction of the population remains within the capture region (Figure 2C). Spatial dispersion *σ*^2^ increases from 9.4 × 10^3^to 2.6 × 10^4^ *μ*m^2^, revealing progressive loss of aggregate compactness and increased fragmentation (Figure 2D).

To identify the governing control parameter, we express the source velocity in dimensionless form as *v*_1_*τ*_run_/*σ*_*s*_, which compares the displacement of the source during a characteristic run time to the spatial scale of the gradient. When plotted against this normalized velocity, tracking success exhibits a clear transition: *η*drops below 0.5 in the range *v*_*f*_ *τ*_run_/*σ*_*s*_ ≈ 0.05–0.10, corresponding to the intermediate regime between *v*_*f*_ = 5and 10 *μ*m/s. Beyond this point, *η*approaches a plateau at low values, indicating a regime shift rather than a gradual degradation of performance.

Taken together, these results demonstrate that dynamic chemotactic tracking is governed by a competition between environmental timescales and intrinsic bacterial response dynamics. When the displacement of the source over a run time remains small compared to the gradient length scale (*v*_*f*_ *τ*_run_ ≪ *σ*_*s*_), populations successfully track the signal. In contrast, when these scales become comparable, the system undergoes a transition to a degraded tracking regime, ultimately leading to complete loss of co-localization. This establishes a fundamental limit on chemotactic tracking imposed by the temporal structure of the environment.

### 3.3 Effect of temporal noise structure on chemotactic robustness

#### Noise class comparison: white, pink, Ornstein–Uhlenbeck

Beyond mean signal amplitude, temporal fluctuations in the perceived chemoattractant gradient impose an additional constraint on chemotactic performance. To isolate the role of temporal structure, we compared three canonical noise classes—white, pink (1/f), and Ornstein–Uhlenbeck (temporally correlated), applied to the sensed signal increment Δm at matched variance.

Spatial snapshots reveal clear qualitative differences across noise classes (Figure 3A–C). Under white noise, populations remain largely confined within the capture region under both static and dynamic conditions, forming compact aggregates centered on the source. Pink noise produces broader spatial distributions and increased lag under dynamic tracking. In contrast, temporally correlated Ornstein– Uhlenbeck noise induces the strongest disruption, leading to elongated, fragmented aggregates and loss of confinement. These qualitative differences are reflected in steady-state metrics (Figure 3D–F). Localization efficiency decreases and spatial dispersion increases progressively from white to pink to Ornstein–Uhlenbeck noise, while aggregate persistence and success probability are strongly reduced in the correlated regime. The temporal evolution of η(t) further shows that white noise produces trajectories nearly indistinguishable from the no-noise case, whereas correlated noise introduces larger fluctuations and lower steady-state performance. Together, these results establish a clear hierarchy of robustness: no noise ≈ white > pink > correlated (Ornstein–Uhlenbeck), demonstrating that temporal correlations—not noise amplitude—are the dominant factor governing chemotactic disruption.

**Figure 3.**
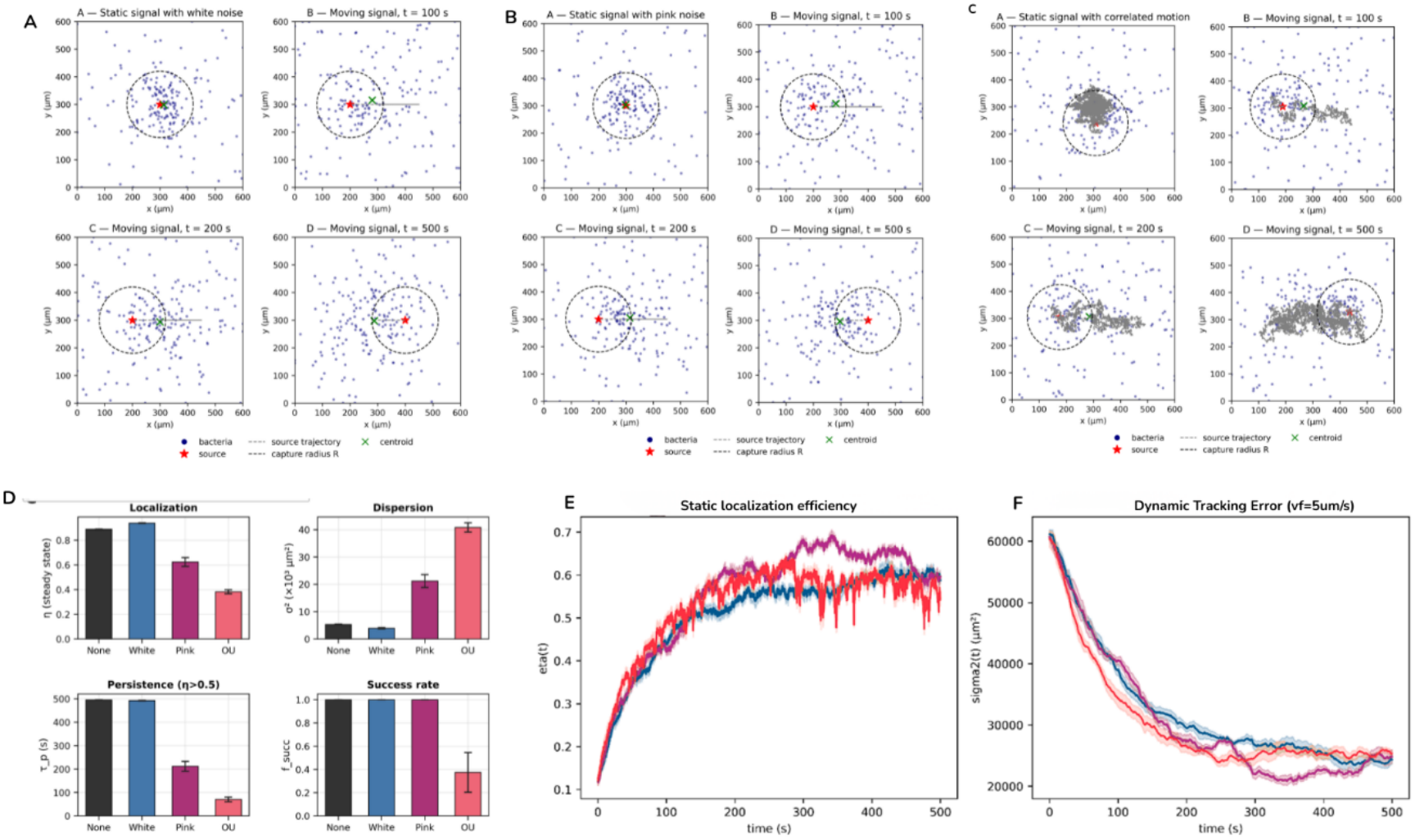
Effect of temporal noise structure on chemotactic robustness. (A–C) Spatial snapshots for white, pink (1/f), and correlated (Ornstein–Uhlenbeck) noise under static and dynamic conditions, showing increasing aggregate dispersion and loss of confinement with temporal correlation. (D) Steady-state metrics of localization, dispersion, persistence, and success rate, revealing a robustness hierarchy from no noise/white to correlated noise. (E) Temporal evolution of localization efficiency η(t), highlighting increased variability and reduced steady-state performance for correlated noise. (F) Evolution of spatial dispersion under dynamic tracking (v_f_ = 5 μm/s), showing greater and more persistent spreading for temporally correlated fluctuations.

#### Correlation-time sweep: identification of a critical sensing timescale

To move beyond discrete noise classes, we next treated temporal correlation as a continuous control parameter. Using the Ornstein–Uhlenbeck process, we systematically varied the correlation time over more than two orders of magnitude (τ_c/τ_run = 0.1–30), while keeping noise amplitude constant. Under static conditions, localization efficiency decreases with increasing normalized correlation time (Figure 4A). For τ_c/τ_run ≪ 1, the system remains robust (η ≈ 0.89), indicating that fast fluctuations are effectively filtered by temporal sensing. As τ_c increases, performance declines sharply, with the steepest drop occurring between τ_c/τ_run ≈ 0.3 and 1.0. Beyond this regime, localization degrades substantially, reaching η ≈ 0.34 at τ_c/τ_run = 10. Aggregate persistence exhibits an even sharper transition (Figure 4B), collapsing from near-complete stability (τ_p ≈ 490 s) to short-lived aggregates (τ_p ≈ 70 s) as τ_c crosses τ_run. In parallel, spatial dispersion increases by nearly an order of magnitude (Figure 4C), indicating that loss of localization corresponds to real spatial spreading of the population. Under dynamic tracking conditions (v_f = 5 μm/s), the system is even more sensitive to temporal correlations (Figure 4D). Tracking error increases and localization decreases, with the transition to degraded behavior occurring slightly earlier (τ_c/τ_run ≈ 0.3–1.0).

**Figure 4.**
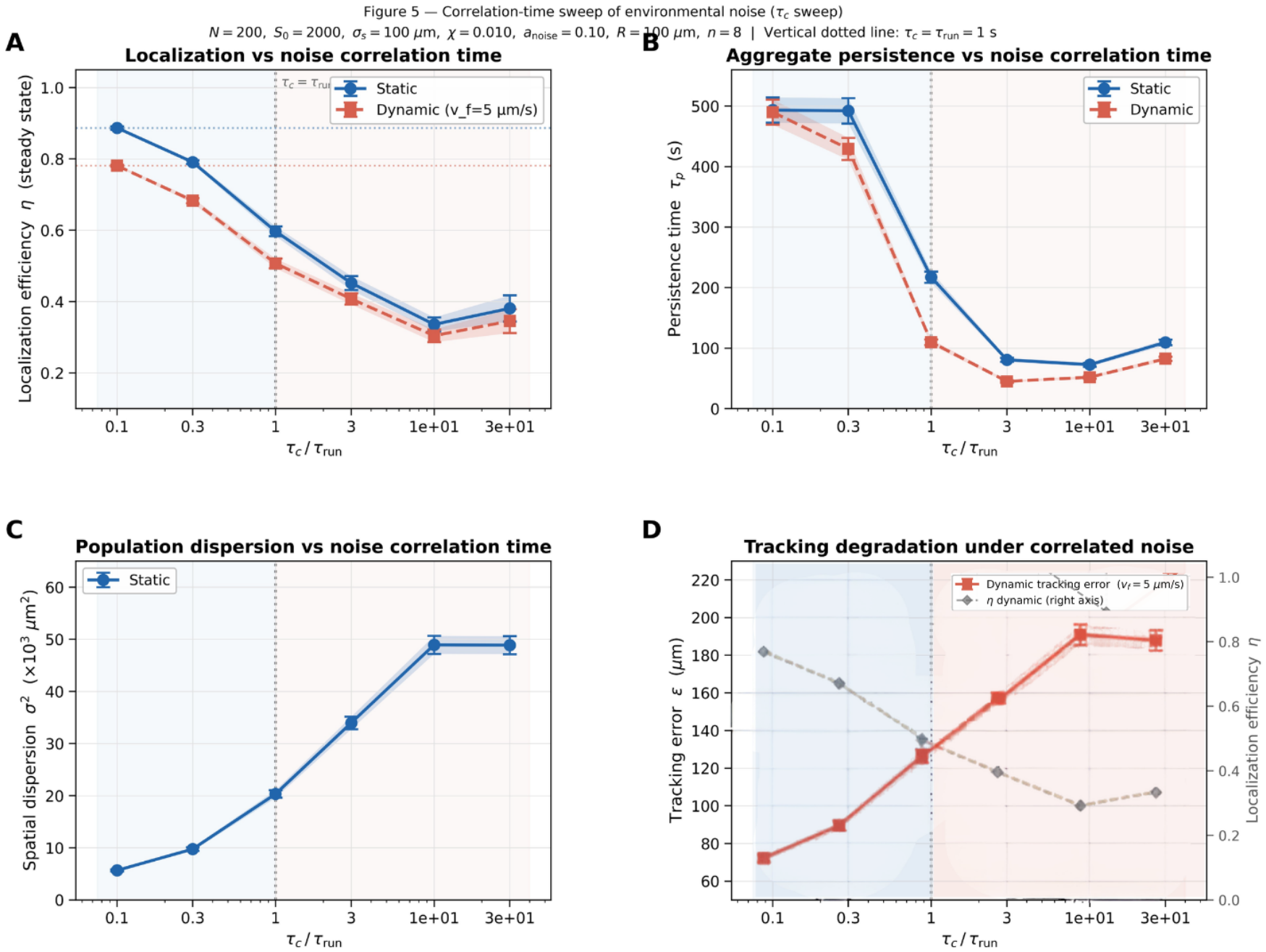
Correlation-time dependence of chemotactic performance. (A) Localization efficiency ηdecreases with increasing normalized correlation time τ_+_/τ_run_, with a sharp transition near τ_c_ ≈ τ_run_. (B) Aggregate persistence τ_p_ collapses across the same threshold, indicating loss of stability under temporally correlated fluctuations. (C) Spatial dispersion σ^1^increases monotonically with τ_c_, reflecting progressive loss of confinement. (D) Under dynamic tracking (v_f_ = 5 μm/s), tracking error εincreases while localization efficiency decreases, showing enhanced sensitivity to correlation time. The vertical dashed line marks τ_c_ = τ_run_, separating regimes of fast (filtered) and slow (disruptive) fluctuations.

These results identify a *critical correlation timescale τ_c ≈ τ_run*\* that separates two regimes: fast fluctuations (τ_c ≪ τ_run): effectively averaged out, and slow fluctuations (τ_c ≳ τ_run): persist over the sensing window and bias gradient estimation. A weak recovery at very large τ_c reflects quasi-stationary noise segments rather than restored robustness. Thus, the ratio τ_c/τ_run emerges as the key dimensionless parameter controlling the transition from robust chemotaxis to noise-induced failure.

#### Spectral analysis: low-frequency power as a predictor of chemotactic failure

The correlation-time sweep shows that slow fluctuations are disproportionately disruptive. This suggests that chemotactic performance is governed not directly by τ_c, but by how noise power is distributed across temporal frequencies relative to the sensing bandwidth. To formalize this, we analyzed the power spectral density (PSD) of the noise signals and defined the low-frequency power fraction:

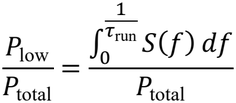

which quantifies the proportion of signal power contained at frequencies slower than the bacterial response timescale.

As τ_c increases, PSDs shift systematically toward lower frequencies (Figure 5A). The corresponding low-frequency fraction increases monotonically from 0.37 to 0.995 and matches the theoretical prediction (Figure 5B), confirming the consistency of the noise implementation. Crucially, this spectral metric provides a strong predictor of chemotactic performance. Localization efficiency decreases approximately linearly with increasing low-frequency content (r = −0.932, p = 0.007; Figure 5C), and aggregate persistence shows a similarly strong negative correlation (r = −0.911, p = 0.011; Figure 5D). In both cases, performance deteriorates rapidly once the low-frequency fraction exceeds ≈ 0.9, consistent with the transition identified in the correlation-time sweep. In contrast, total signal variance remains constant across all conditions and exhibits no predictive power (Figure 5E). Systems with identical variance but different spectral composition display markedly different outcomes, demonstrating that amplitude alone is insufficient to characterize environmental perturbations.

**Figure 5.**
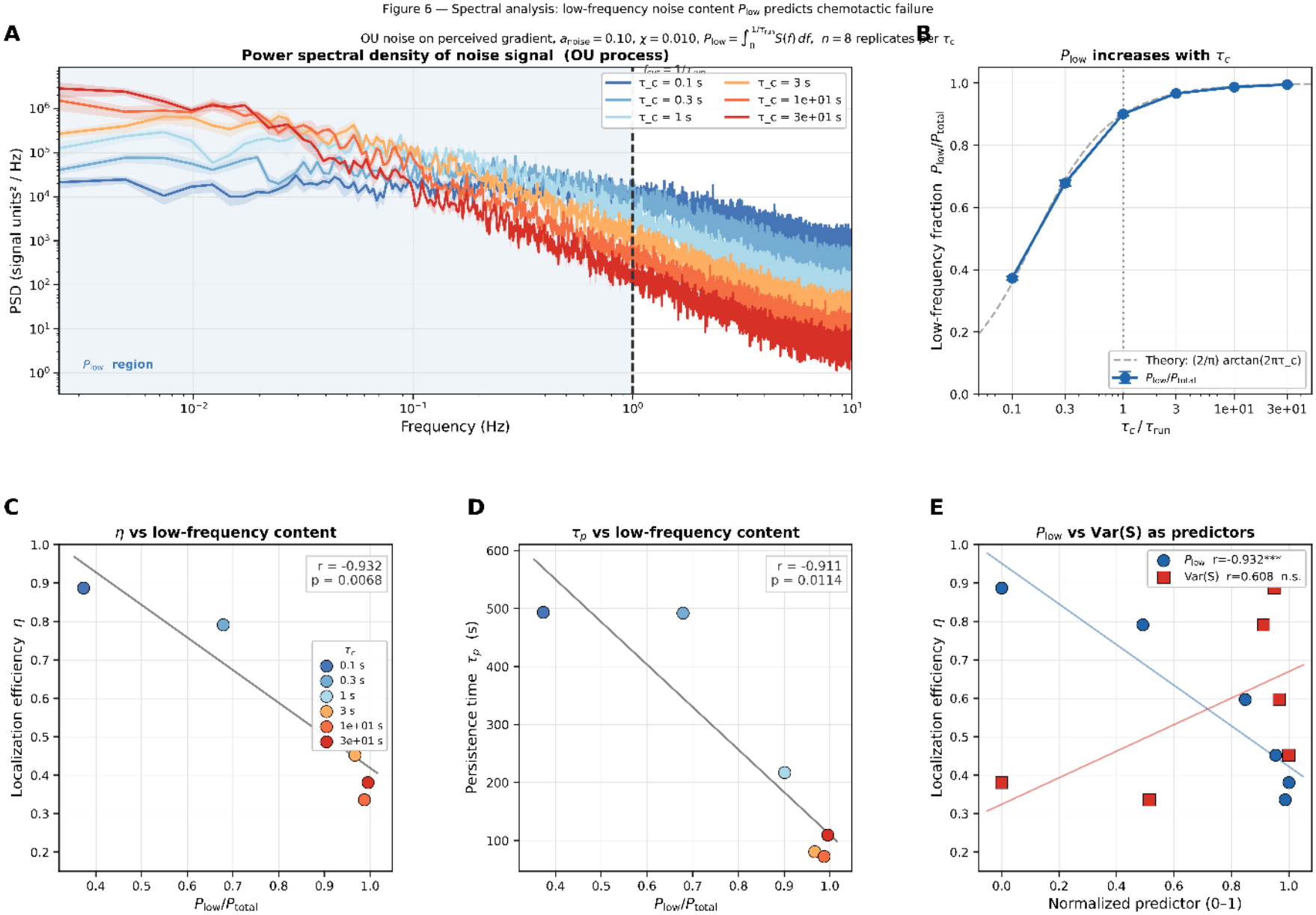
Spectral predictor of chemotactic failure. (A) Power spectral densities (PSDs) of Ornstein–Uhlenbeck (OU) noise for increasing correlation times τ_c_, showing a systematic shift of power toward low frequencies; shaded region indicates P_low_ below 1/τ_run_ (B) Low-frequency power fraction P_low_/P_total_ increases monotonically with τ_c_, matching theoretical predictions. (C–D) Localization efficiency ηand aggregate persistence τ_p_ decrease strongly with increasing low-frequency content, demonstrating that performance degrades as slow fluctuations dominate. (E) Comparison of predictors shows that P_low_/P_total_ strongly correlates with failure, whereas total variance Var(S) has no predictive power.

These results establish a general principle: chemotactic populations act as temporal filters, suppressing high-frequency fluctuations while remaining sensitive to low-frequency components. As a consequence, the fraction of low-frequency power—not total variance—determines whether fluctuations are averaged out or persistently bias gradient sensing. The low-frequency power fraction therefore provides a unified predictor of both localization failure and tracking breakdown, linking environmental noise statistics directly to emergent population-level behavior.

### 3.4 Flow-Induced Destabilization

To prove the robustness of chemotactic aggregates to external transport, we introduced vertical flow fields after aggregates had reached steady state under static signaling conditions (*η*_pre_ ≈ 0.42). Flow was activated at *t* = 150s, and its effect on aggregate stability was quantified as a function of geometry, intensity, and topology.

A schematic of the flow configuration is shown in Figure 6A. Flow was imposed in vertical strips positioned symmetrically or asymmetrically relative to the signal center, with the key control parameter being the distance *d*between the inner edge of the flow region and the center of the chemotactic capture region *R*. This setup allows direct control over whether the imposed flow overlaps with the region where chemotactic drift is sufficient to retain cells. Spatial snapshots following flow activation reveal a clear qualitative transition (Figure 6B). When flow is confined far outside the capture region (*d* ≥ 150–200 *μ*m), the aggregate remains compact and centered on the source. At intermediate distances (*d* ≈ 100 *μ*m), corresponding to the boundary of *R*, the aggregate is only weakly perturbed. In contrast, when flow penetrates the capture region (*d* = 50 *μ*m), the aggregate disperses rapidly, with a marked reduction in density and visible loss of coherence. This transition is quantified by the aggregate persistence time *τ*_*p*_ (Figure 6C). For flow positioned outside the capture region, post-flow persistence remained maximal within the observation window (*τ*_*p*_ ≈ 110 s) across all tested flow speeds. However, when the inner edge of the flow strip enters *R*, persistence collapses sharply, dropping to *τ*_*p*_ ≈ 82s at *u*_*f*_ = 10 *μ*m/sand decreasing further with increasing flow intensity. The transition occurs at *d* ≈ *R*, demonstrating that the onset of destabilization is governed by geometric overlap rather than flow strength alone. Flow intensity modulates the severity of this transition once overlap is established. At the boundary case (*d* = 100 *μ*m), increasing flow speed produces a monotonic reduction in post-flow localization and persistence. For example, *η*_post_ remains close to baseline at low velocities but decreases substantially at higher speeds (*u*_*f*_ = 20–40 *μ*m/s), accompanied by a reduction in persistence time. This indicates that while geometry sets the threshold for destabilization, intensity controls the rate and extent of aggregate breakdown.

**Figure 6.**
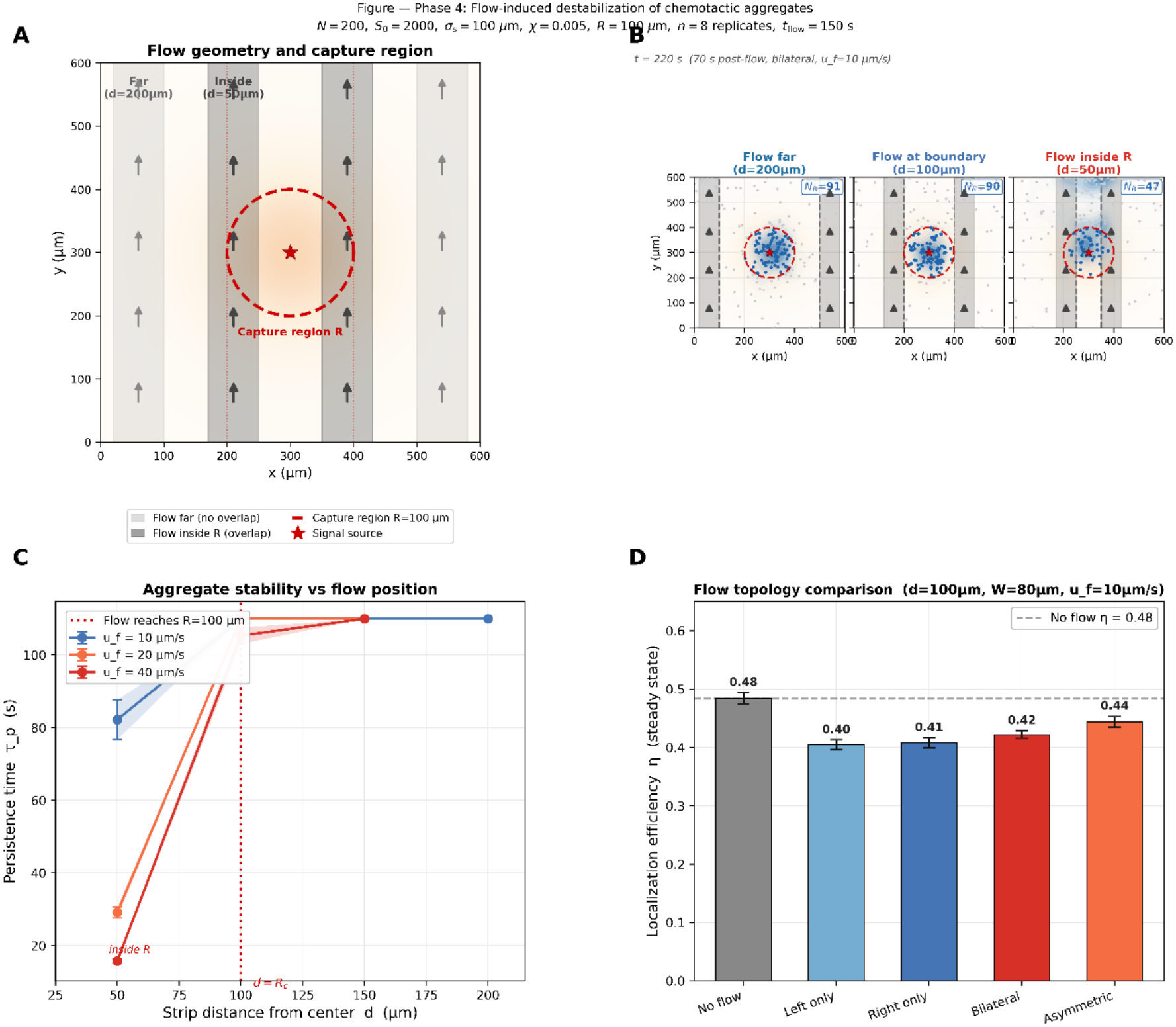
Flow-induced destabilization of chemotactic aggregates. (A) Schematic of flow geometry relative to the chemotactic capture region R, showing configurations with flow outside, at the boundary, and inside the aggregate core. (B) Spatial snapshots 70 s after flow onset (u_f_ = 10 μm/s), illustrating progressive aggregate disruption as flow penetrates the capture region. (C) Aggregate persistence τ_p_ as a function of strip distance d, revealing a sharp transition in stability when flow overlaps with R. (D) Comparison of flow topologies at marginal overlap (d = 100 μm), showing that asymmetric configurations preserve higher localization by maintaining a recovery pathway.

The role of spatial extent was examined by varying the strip width *W*at fixed position. Changes in width produced comparatively minor effects relative to those induced by shifting the inner edge position. Even narrow strips caused substantial destabilization when they intersected the capture region, whereas increasing width outside *R*had little additional impact. This demonstrates that the critical factor is the presence of flow within the high-gradient core, rather than the total spatial extent of the perturbation. Flow topology further modulates aggregate stability (Figure 6D). At marginal overlap (*d* = 100 *μ*m, *u*_*f*_ = 10 *μ*m/s), unilateral and symmetric bilateral flows produce similar reductions in localization efficiency. However, asymmetric bilateral configurations consistently preserve higher localization, indicating enhanced robustness. This effect becomes more pronounced at higher flow speeds, where symmetric configurations lead to stronger depletion.

The enhanced stability under asymmetric flow can be attributed to the presence of a weakly perturbed flank that acts as a reservoir of cells. In such configurations, bacteria displaced from the high-flow region can re-enter the capture region via chemotactic drift from the less perturbed side. In contrast, symmetric flows impose comparable advective removal on both sides, eliminating recovery pathways and leading to sustained depletion.

Taken together, these results demonstrate that the stability of chemotactic aggregates under flow is governed primarily by the spatial overlap between the flow field and the capture region. Flow confined outside *R*is effectively inert, whereas flow penetrating *R*induces a sharp transition to destabilization. The severity of this transition is then modulated by flow intensity and topology, reflecting the balance between advective transport and chemotactic retention. This establishes a general principle: external transport disrupts collective chemotaxis only when it operates within the region where temporal sensing can sustain aggregation.

## 4. Discussion

### 4.1 Minimal mechanisms for emergent chemotactic organization

The results presented here demonstrate that robust collective organization can emerge from minimal behavioral rules, without explicit inter-agent interactions. In our framework, each agent relies solely on local temporal sensing of chemoattractant signals, yet the population exhibits coherent aggregation, adaptive tracking, and structured responses to environmental perturbations. This contrasts with many active matter models that require alignment or explicit coupling, highlighting instead the central role of the spatiotemporal structure of the environment as the primary organizing driver. This behavior is consistent with the canonical chemotactic strategy of *Escherichia coli*, in which cells bias run durations based on temporal signal variation rather than direct coordination. Our results extend this principle to the population level, showing that such minimal sensing rules are sufficient to generate large-scale organization, provided that environmental dynamics remain within the response bandwidth of the chemotactic mechanism.

### 4.2 Dynamic limits of chemotactic tracking

A central result of this study is the identification of a velocity-dependent limit in chemotactic tracking. As the speed of the signal source increases, populations transition from coherent tracking to fragmentation and ultimately to loss of co-localization. This transition reflects a mismatch between environmental dynamics and the intrinsic response timescale of the chemotactic mechanism. Because cells rely on temporal comparisons, rapid displacement of the signal reduces the reliability of directional inference, leading to increased tracking error and delayed reorientation. Importantly, this limitation arises even in the absence of noise, establishing a fundamental upper bound on controllability imposed purely by dynamics. This behavior can be interpreted in terms of an effective chemotactic capture region: the spatial domain within which chemotactic drift can maintain localization against stochastic motion. When signal dynamics cause cells to exit this region faster than they can reorient, tracking fails.

### 4.3 Temporal noise and collective temporal filtering

Beyond deterministic dynamics, we show that the temporal structure of environmental noise plays a dominant role in shaping collective behavior. Populations respond qualitatively differently to white, pink, and correlated noise, even at identical amplitudes. High-frequency fluctuations are largely averaged out by temporal sensing, whereas slow, correlated fluctuations persist over timescales comparable to the cellular response and introduce systematic errors in gradient estimation. This asymmetry reveals that chemotactic populations act as temporal filters, suppressing fast fluctuations while remaining sensitive to slow variations. This constitutes a key conceptual contribution: temporal correlations—not noise amplitude—emerge as the primary control parameter governing robustness. This extends previous work focused on uncorrelated or intracellular noise by explicitly linking environmental temporal structure to population-level behavior.

### 4.4 Critical timescale and spectral criterion for failure

Building on this filtering mechanism, we identify a quantitative threshold separating robust from failed chemotactic behavior. The correlation-time sweep reveals a critical regime at

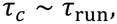

where performance degrades sharply. When fluctuations are faster than the sensing timescale, they are effectively averaged out; when they persist longer, they bias navigation. This result admits a more general formulation in spectral terms. We show that performance is controlled by the fraction of signal power at frequencies below 1/*τ*_run_ rather than by total variance. The low-frequency power fraction provides a strong predictor of both localization and tracking failure, whereas total signal variance has no predictive value. Together, these findings establish a unified principle: chemotactic breakdown occurs when environmental fluctuations contain sufficient low-frequency power to evade temporal averaging. This provides a population-level analogue of sensing limits and links environmental statistics directly to emergent behavior.

### 4.5 Flow-induced breakdown and spatial control of robustness

External transport introduces an additional mechanism of disruption that is fundamentally spatial rather than temporal. We show that flow-induced destabilization is governed not by flow magnitude alone, but by its spatial overlap with the chemotactic capture region. Flow confined outside this region has negligible effect, whereas flow penetrating the high-gradient core induces a sharp transition to aggregate collapse. Once overlap is established, flow intensity controls the rate of destabilization, while topology determines the availability of recovery pathways. In particular, asymmetric configurations preserve partially protected regions that act as reservoirs, enabling re-entry via chemotactic drift. This establishes a second key principle: chemotactic robustness under flow is determined by geometric overlap between transport and the sensing-active region.

### 4.6 Unified robustness framework

Taken together, these results define a unified phase-space of chemotactic robustness governed by three primary control parameters: normalized source velocity *v*_1_*τ*_run_/*σ*_*s*_ , normalized correlation time *τ*_0_/*τ*_run_, and spatial overlap between perturbations and the capture region. These parameters define distinct regimes: a robust regime with stable aggregation and tracking, an intermediate regime with partial fragmentation, and a failure regime characterized by loss of organization. Importantly, temporal noise and flow act synergistically. Slowly varying fluctuations effectively displace the signal over extended timescales, increasing the likelihood of escape from the capture region, while flow imposes directional transport that amplifies this effect. Together, they accelerate breakdown beyond what either mechanism produces independently.

### 4.7 Implications and outlook

The framework developed here is broadly applicable across microbiology, active matter, and synthetic systems. Many natural environments—such as tissues, porous media, and microfluidic flows—exhibit both temporal variability and spatially structured transport. Our results provide a quantitative basis for predicting when chemotactic populations can maintain directed behavior versus when they will disperse. From an engineering perspective, these findings define design principles for controlling chemotactic systems. Reliable targeting requires that environmental dynamics remain within the temporal sensing bandwidth and that external transport does not disrupt the capture region. Conversely, controlled disruption can be achieved by introducing low-frequency fluctuations or flow fields that overlap this region. Several limitations remain. The present model neglects inter-agent interactions and detailed intracellular signaling, both of which may modulate robustness at higher densities or in real biological systems. Future work should incorporate these effects and explore experimental validation using microfluidic platforms capable of imposing controlled noise spectra and flow geometries.

